# DNA damage inhibits root growth by enhancing cytokinin biosynthesis in *Arabidopsis thaliana*

**DOI:** 10.1101/2020.06.19.160838

**Authors:** Naoki Takahashi, Soichi Inagaki, Kohei Nishimura, Hitoshi Sakakibara, Ioanna Antoniadi, Michal Karady, Karin Ljung, Masaaki Umeda

## Abstract

Plant root growth is influenced by external factors to adapt to changing environmental conditions. However, the mechanisms by which environmental stresses affect root growth remain elusive. Here we found that DNA double-strand breaks (DSBs) induce the expression of genes for the synthesis of cytokinin hormones and enhance the accumulation of cytokinins in the *Arabidopsis* root tip. This is a programmed response to DSBs through the DNA damage signaling pathway. Our data showed that activation of cytokinin signalling suppresses the expression of *PIN-FORMED* genes that encode efflux carriers of another plant hormone, auxin, thereby disturbing downward auxin flow and causing cell cycle retardation in the G2 phase. Elevated cytokinin signalling also promotes an early transition from cell division to endoreplication, resulting in a reduction of the root meristem size. We propose that in response to DNA stress, plants inhibit root growth by orchestrating hormone biosynthesis and signalling.

---

Plant roots play a crucial role in water and nutrient uptake, anchorage to soil, and sensing the rhizosphere environment^1^. Since these functions have a substantial impact on overall plant growth, root development is precisely controlled in response to changing underground conditions. Environmental stresses usually inhibit root growth; salinity, oxidation or heat stress severely retards root growth. Previous studies demonstrated that high boron or aluminium stress, which causes DNA damage, leads to delay or cessation of cell division, thereby suppressing root growth^2,3^. This is an active response to DNA damage that is governed by the cell cycle checkpoint mechanism, in which cell cycle progression is arrested at a specific stage to ensure DNA repair or to provoke cell death in severe cases^4^. As do other eukaryotes, plants possess two protein kinases, ATAXIA TELANGIECTASIA MUTATED (ATM) and ATM AND RAD3-RELATED (ATR), that sense DNA damage and trigger cell cycle checkpoints^5,6^. ATM is activated by DNA double-strand breaks (DSBs), whereas ATR primarily senses single-strand DNA and replication stress caused by DNA replication fork blocking. In animals and fungi, DNA damage signals are transmitted to Checkpoint-1 (CHK1) and CHK2 kinases, and ATM, ATR, CHK1 and CHK2 phosphorylate and activate the tumour suppressor protein p53^7^. However, orthologues of CHK and p53 are missing in plants; instead, the plant-specific NAC-type transcription factor, named SUPPRESSOR OF GAMMA RESPONSE 1 (SOG1), plays a central role in transmitting the signal from ATM and ATR^8^. SOG1 is phosphorylated and activated by ATM and ATR^9,10^, and binds to the sequence CTT[N]_7_AAG to induce the expression of target genes involved in DNA repair, cell cycle arrest and stem cell death^11,12^.

In *Arabidopsis* roots, cells actively divide and proliferate in the meristematic zone (MZ) in the tip region. After several rounds of cell division, cells stop dividing and start endoreplication, in which DNA replication is repeated without mitosis or cytokinesis. Endoreplicating cells begin to elongate and constitute the transition zone (TZ), and eventually start more rapid cell elongation through actin reorganization in the elongation/differentiation zone (EDZ)^13,14^. It was shown that DSBs arrest the cell cycle at G2 in the MZ^15^ and promote an early onset of endoreplication^16^, reducing the meristem size and inhibiting root growth. These DNA damage responses (DDRs) are under the control of the ATM-SOG1 pathway^15,16^. Our previous study showed that two repressor-type R1R2R3-Myb transcription factors (Rep-MYBs), MYB3R3 and MYB3R5, participate in DNA damage-induced G2 arrest by suppressing the expression of G2/M-specific genes^15^. We have recently demonstrated that protein accumulation of Rep-MYBs is regulated by the transcription factors ANAC044 and ANAC085^17^, while cyclin-dependent kinase (CDK) also plays a crucial role in Rep-MYB accumulation; namely, Rep-MYBs phosphorylated by CDKs are targeted for degradation, whereas low CDK activity caused by DNA damage stabilizes Rep-MYB proteins, leading to G2 arrest^15^. However, it remains unclear how CDK activities are inhibited by DNA stress in the root tip. Although DNA damage induces the expression of several CDK inhibitors and represses the gene encoding activator-type R1R2R3-Myb transcription factor (Act-MYB), which induces G2/M-specific genes, such transcriptional responses are not sufficient to cause G2 arrest^12,15,18^. Therefore, some unknown mechanism(s) should function in downregulation of CDK activities under DNA stress.

Cytokinin plant hormones participate in various developmental and physiological processes such as seed germination, flowering and senescence^19^. The initial step of cytokinin biosynthesis is catalysed by ATP/ADP isopentenyltransferases (IPTs), which produce ribosylated and phosphorylated precursors of N^6^-(Δ^2^-isopentenyl) adenine (iP)^20,21^. These products are converted to ribosylated and phosphorylated forms of *trans*-zeatin (*t*Z) by the action of CYP735A, which hydroxylates the *trans*-end of the prenyl side chain^22,23^. LONELY GUYs (LOGs) are essential enzymes for converting those precursors to biologically active cytokinins^24^. Previous studies demonstrated that cytokinins are systemically transported in *Arabidopsis*: iP-types are moved from shoots to the root meristem via the phloem^25,26,27^, and *t*Z-types are transported from roots to shoots via the xylem^28^. CYP735A2 is predominantly expressed in developing vascular tissues of roots, thereby converting iP- to *t*Z-type and supplying *t*Z-types to shoots^28^. The ABC transporter ABCG14 contributes to *t*Z-type cytokinin transport by facilitating xylem loading in roots^29,30^.

In *Arabidopsis*, iP and *t*Z are perceived by three receptors, ARABIDOPSIS HISTIDINE KINASE 2 (AHK2), AHK3 and AHK4/CRE1^31–33^. The cytokinin signal activates the transcription factors, type-B ARABIDOPSIS RESPONSE REGULATORs (ARRs), via the His-Asp phosphorelay pathway^34^. In roots, cytokinin-activated ARR2, which accumulates around the TZ, induces the expression of *CCS52A1* encoding an activator of the E3 ubiquitin ligase, anaphase-promoting complex/cyclosome (APC/C)^35^. This promotes degradation of mitotic regulators such as cyclins, enhancing the transition from the mitotic cell cycle to endoreplication. It was reported that ARR1 and ARR12 also accumulate around the TZ and induce *SHORT HYPOCOTYL 2* (*SHY2*), which encodes a member of the Aux/IAA protein family of auxin signalling repressors^36^. This induction represses the expression of the auxin efflux carrier *PIN-FORMED* (*PIN*) and restricts meristem size. However, the relationship between suppression of *PIN* expression and inhibition of cell division has not yet been carefully studied.

As described above, DNA damage causes G2 arrest and an early onset of endoreplication, and cytokinins function in restricting root meristem size in the absence of external stress. These facts prompted us to hypothesize that cytokinins are associated with DDR in roots. In this study, we found that DSBs induce several cytokinin biosynthesis genes and increase the endogenous cytokinin level in *Arabidopsis* roots, thereby suppressing the expression of *PIN1* and *PIN4* and decreasing the auxin level in the MZ. We propose that enhanced cytokinin biosynthesis is a key to inhibit cell division and promote an early onset of endoreplication in response to DNA damage.

## Results

### DSBs activate cytokinin signalling in roots

To test whether cytokinin signalling is affected by DNA damage in roots, we first examined the promoter activity of the cytokinin-inducible type-A response regulator gene *ARR5*^37^. Five-day-old *Arabidopsis* seedlings were transferred onto a medium containing 8 μM DSB-inducing reagent zeocin^16^, and subjected to GUS staining after 24 h. GUS activity was increased in the vasculature around the boundary between the MZ and the TZ, vascular initial cells and their daughters, and the root cap (Fig. 1a; Supplementary Fig. 1). We also observed roots carrying the cytokinin response marker *Two Component Signaling Sensor new* (*TCSn*)*:GFP*, which reflects the transcriptional activity of type-B response regulators^38^. The expression pattern of this marker was similar to that of *pARR5:GUS*, except that GFP fluorescence was also detected in the epidermis (Fig. 1b). Quantification of GFP fluorescence revealed that the expression level was elevated by zeocin treatment in four epidermal cells above and below the boundary between the MZ and the TZ (Fig. 1c). A higher level of GFP fluorescence was also observed in vascular cells encompassing a 100-μm region around the boundary (Fig. 1d). To confirm that enhanced cytokinin signalling is not a consequence of tissue injury caused by DNA damage, we examined the expression patterns of several cell type-specific markers, *pAHP6:GFP* (protoxylem), *pAPL:GFPer* (phloem), *pCO2:H2B-YFP* (cortex), *pSCR:GFP-SCR* (endodermis and QC) and *pWOX5:NLS-YFP* (QC). As shown in Supplementary Fig. 2, zeocin treatment did not change the expression pattern of any marker, indicating that our experimental condition did not cause severe injury in root tissues. We found that increased expression of *TCSn:GFP* was also observed after 1.5 mM aluminium treatment for 24 h, which is also known to cause DSBs^39^ (Supplementary Fig. 3).

**Figure 1.**
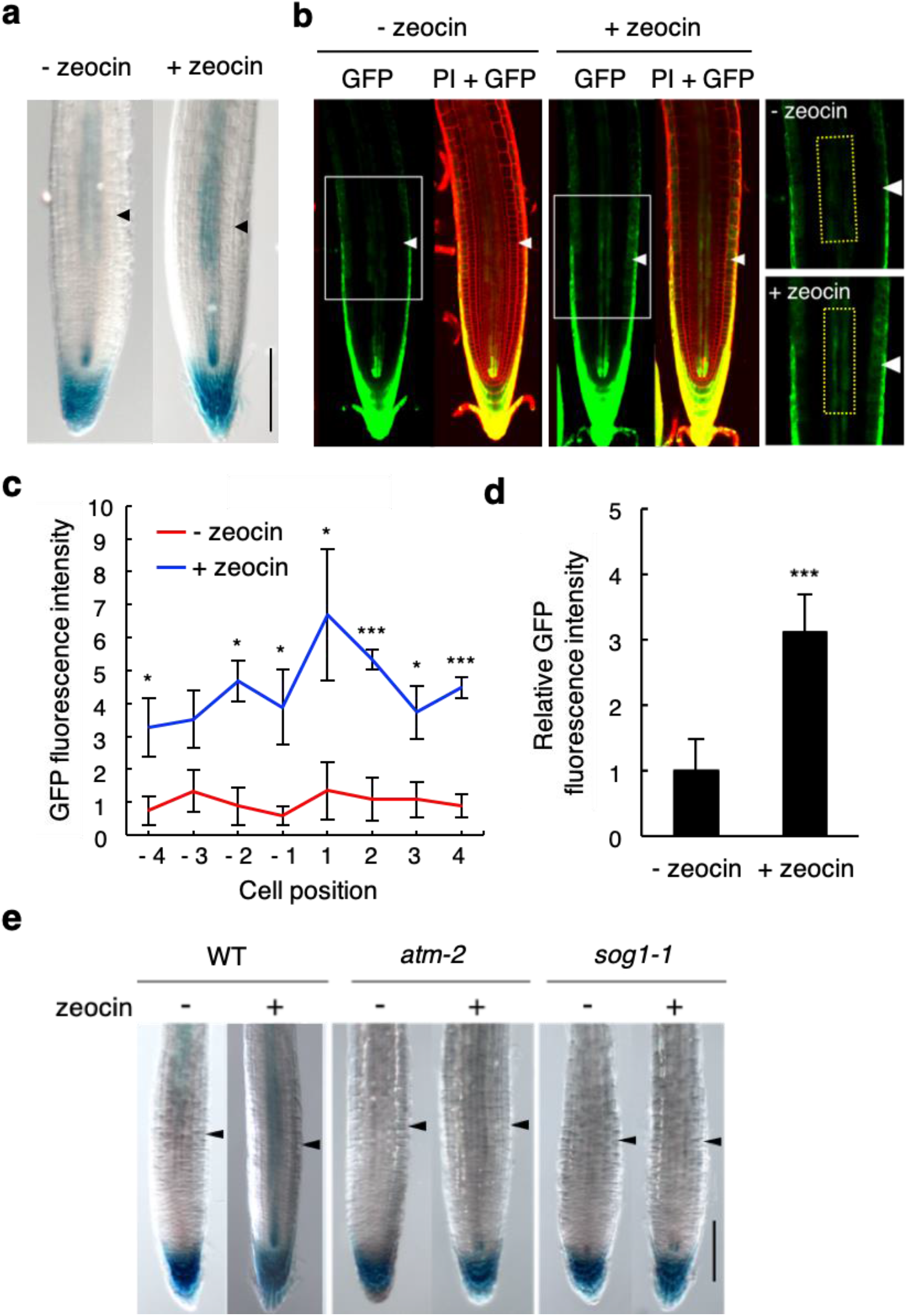
DSBs activate cytokinin signalling in the root tip. (**a**) Zeocin response of the *ARR5* promoter activity. Five-day-old *pARR5-GUS* seedlings were transferred to MS plates supplemented with (+ zeocin) or without (− zeocin) 8 μM zeocin, and grown for 24 h, followed by GUS staining. Arrowheads indicate the boundary between the MZ and the TZ. Bar = 100 μm. (**b**) Zeocin response of the synthetic cytokinin reporter *TCSn:GFP*. Five-day-old *TCSn:GFP* seedlings were treated with or without 8 μM zeocin for 24 h, and GFP fluorescence was observed after counterstaining with PI. Arrowheads indicate the boundary between the MZ and the TZ. Magnified images of the areas marked by white boxes are shown on the right (see also (**d**)). Bars = 100 μm. (**c**) GFP fluorescence intensity in the root epidermis of *TCSn:GFP* seedlings. Cell position ‘1’ indicates the first endoreplicated cell, which is preceded by the last mitotic cell before entry into the endoreplication (cell position ‘-1’). Data are presented as mean ± SD (n > 8). Significant differences from the control without zeocin treatment were determined by Student’s *t*‐test: *, *P* < 0.05; ***, *P* < 0.001. (**d**) GFP fluorescence intensity in the vasculature of *TCSn:GFP* seedlings. GFP fluorescence was measured in the areas surrounded by yellow dotted lines shown on the right in (**b**), which encompass a 100-μm region around the boundary between the MZ and the TZ. The value relative to that of the control without zeocin treatment is shown. Data are presented as mean ± SD (n = 10). The significant difference from the control was determined by Student’s *t*‐test: ***, *P* < 0.001. (**e**) *ARR5* promoter activities in *atm* and *sog1*. Five-day-old seedlings of wild-type (WT), *atm-2* and *sog1-1* harbouring *pARR5:GUS* were transferred to MS plates with (+) or without (−) 8 μM zeocin, and grown for 24 h, followed by GUS staining. Arrowheads indicate the boundary between the MZ and the TZ. Bar = 100 μm.

In response to DSBs, ATM phosphorylates and activates the plant-specific transcription factor SOG1, which then induces hundreds of genes to trigger DDR^8,9^. We found that zeocin treatment increased *pARR5:GUS* expression in wild-type, but not in the *atm-2* or *sog1-1* knockout mutant (Fig. 1e; Supplementary Fig. 1). This indicates that activation of cytokinin signalling in roots is a programmed response to DSBs that requires the ATM–SOG1 pathway.

### DSBs elevate cytokinin level in the root tip

Previous studies demonstrated that the type-B response regulators *ARR1* and *ARR2* are expressed around the TZ and upregulate cytokinin signalling^35,40^. However, our quantitative real-time PCR (qRT-PCR) data showed that neither *ARR1* nor *ARR2* was induced by zeocin in roots (Supplementary Fig. 4a). The marker lines *pARR1:ARR1–GUS* and *pARR2:ARR2–GUS* displayed no change in expression levels after zeocin treatment (Supplementary Fig. 4b). These results indicate that *ARR1* and *ARR2* are not associated with DSB-dependent activation of cytokinin signalling.

We therefore examined whether the endogenous cytokinin level increases in response to DNA damage. Cytokinin contents were separately measured in cells constituting the TZ or the MZ. We performed fluorescence-activated cell sorting (FACS) on GFP- or CFP-expressing protoplasts prepared from *ROOT CLAVATA HOMOLOG 1* (*RCH1*) *promoter:GFP* or *RCH2 promoter:CFP* reporter lines, respectively. Since the *RCH1* and *RCH2* promoters are active in the MZ and the TZ^40^, respectively, isolated protoplasts are expected to be derived from each zone^41^. Our cytokinin measurements revealed that in the presence of zeocin, the levels of *t*Z, and cytokinin glucosides *t*Z-9-*N*-glucoside (*t*Z9G) and isopentenyladenosine-7-*N*-glucoside (iP7G) increased in the TZ, but not in the MZ (Fig. 2). The amounts of cytokinin precursors *trans-*zeatin-riboside (*t*ZR) and isopentenyladenosine-riboside (iPR), and cytokinin glucosides *t*Z-*O*-glucoside (*t*ZOG) and *t*Z-7-*N*-glucoside (*t*Z7G), were elevated in both the MZ and the TZ, although the increases were higher in the TZ (Fig. 2). These results suggest that DSBs increase the endogenous cytokinin level in the MZ and more highly in the TZ.

**Figure 2.**
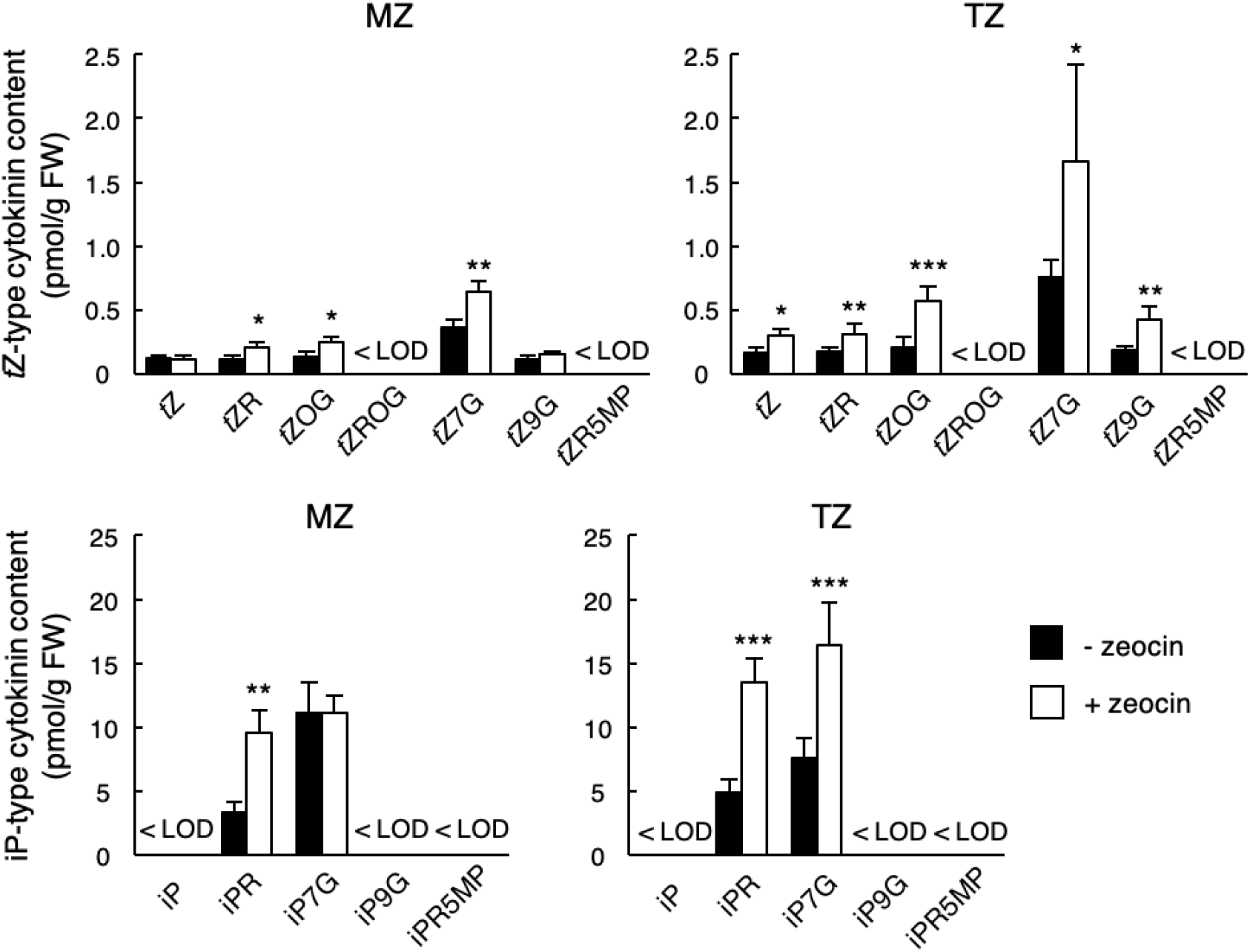
DSBs elevate cytokinin level in the root tip. Five-day-old seedlings of the marker lines *pRCH1:GFP* and *pRCH2:CFP* were transferred to MS plates with (+ zeocin) or without (− zeocin) 8 μM zeocin, and grown for 24 h. FACS was conducted to collect GFP- or CFP-positive protoplasts, which were analysed for their cytokinin concentration using LC-MS/MS. Amounts of iP- and *t*Z-type cytokinins per gram of fresh weight (FW) in GFP- or CFP-positive protoplasts, which were derived from cells in the MZ or the TZ, respectively, are shown. Data are presented as mean ± SD (n = 4). Significant differences from the control without zeocin treatment were determined by Student’s *t*-test: *, *P* < 0.05; **, *P* < 0.01; ***, *P* < 0.001. < LOD, below limit of detection. *t*Z, *trans*-zeatin; *t*ZR, *t*Z riboside; *t*ZOG, *t*Z-O-glucoside; *t*ZROG, *t*ZR-O-glucoside; *t*Z7G, *t*Z-7-glucoside; *t*Z9G, *t*Z-9-glucoside; *t*ZR5MP, *t*ZR-5-monophosphate; iP, *N*^6^-(Δ^2^-isopentenyl) adenine; iPR, iP riboside; iP7G, iP-7-glucoside; iP9G, iP-9-glucoside; iPR5MP, iPR-5-monophosphate.

### DSB-dependent induction of cytokinin biosynthesis genes inhibits root growth

To identify the cause of DNA damage-induced cytokinin accumulation, we measured the transcript levels of 17 cytokinin biosynthesis genes: seven ATP/ADP *IPT*s (*IPT1* and *IPT3*– *IPT8*), two *CYP735A*s (*CYP735A1* and *CYP735A2*) and eight *LOG*s (*LOG1*–*LOG8*). We did not analyse tRNA *IPT*s (*IPT2* and *IPT9*) because they are engaged in the synthesis of *cis*-zeatin (*c*Z), a less physiologically active cytokinin than iP or *t*Z^42^. qRT-PCR using RNA from whole seedlings revealed that zeocin treatment elevated the mRNA levels of *IPT1*, *IPT3*, *IPT5*, *IPT7*, *CYP735A2* and *LOG7*, but not *LOG3*, *LOG4* or *LOG8* (Fig. 3a). We could not detect transcripts of *IPT4*, *IPT6*, *IPT8*, *CYP735A1*, *LOG1*, *LOG2*, *LOG5* or *LOG6* regardless of zeocin treatment, probably due to very low expression. Induction of *IPT1*, *IPT3*, *IPT5*, *IPT7*, *CYP735A2* and *LOG7* was not observed in the *atm-2* or *sog1-1* mutant (Supplementary Fig. 5), indicating that their induction is under the control of the ATM–SOG1 pathway.

**Figure 3.**
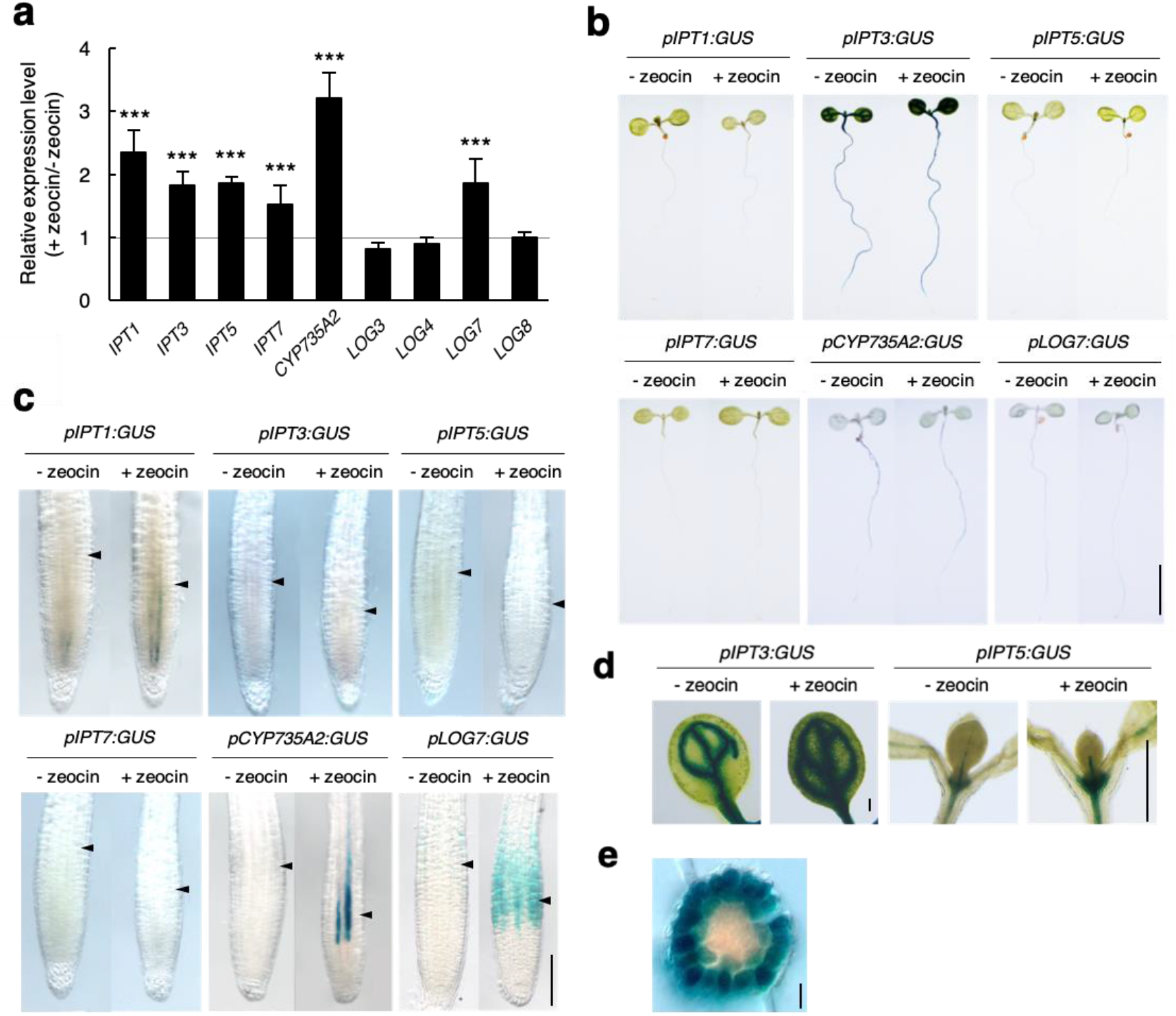
DSBs induce cytokinin biosynthesis genes. (**a**) Transcript levels of cytokinin biosynthesis genes. Five-day-old wild-type seedlings were transferred to MS plates with or without 8 μM zeocin, and grown for 24 h. Total RNA was extracted from whole seedlings and subjected to qRT-PCR. Transcript levels of cytokinin biosynthesis genes were normalized to that of *ACTIN2*, and are indicated as relative values, with that of the control without zeocin treatment set to 1. Data are presented as mean ± SD calculated from three biological and technical replicates. Significant differences from the control were determined by Student’s *t*-test: ***, *P* < 0.001. (**b–e**) Zeocin response of *pIPT1:GUS*, *pIPT3:GUS, pIPT5:GUS*, *pIPT7:GUS*, *pCYP735A2:GUS* and *pLOG7:GUS*. Five-day-old seedlings were treated with (+ zeocin) or without (− zeocin) 8 μM zeocin for 24 h. GUS-stained samples were observed for whole seedlings (**b**), root tips (**c**) and cotyledons and shoot apices (**d**). The cross section around the TZ of zeocin-treated *pLOG7:GUS* is shown in (**e**). Arrowheads in (**c**) indicate the boundary between the MZ and the TZ. Bars = 1 cm (**b**), 100 μm (**c**), 1 mm (**d**) and 20 μm (**e**).

Our observation of *promoter:GUS* reporter lines supported the above result. *IPT1*, which is known to be expressed in the procambium^43^, displayed increased expression in zeocin-treated root tips (Fig. 3c). The promoter activities of *IPT3* and *IPT5* were elevated in cotyledons and shoot apices, respectively, but not detected in the root tip (Fig. 3b–d). *pIPT7:GUS* showed no GUS signal regardless of zeocin treatment (Fig. 3b, c), probably because the promoter region used for the reporter construction lacks essential *cis*-element(s). Interestingly, zeocin highly induced *CYP735A2* and *LOG7* in the vasculature and in the epidermis and cortex, respectively, around the TZ (Fig. 3c, e). These results suggest that under DNA damage conditions, induction of *IPT3* and *IPT5* in shoots provides more cytokinin precursors, and that induction of *CYP735A2* and *LOG7* around the TZ leads to higher accumulation of iP- and *t*Z-type cytokinins in the root tip, as revealed by our cytokinin measurement (Fig. 2). The induction of *pLOG7:GUS* was also observed under DSB-causing aluminium stress (Supplementary Fig. 6).

We then examined whether the induction of cytokinin biosynthesis genes is involved in root growth inhibition in response to DNA damage. Since the *ipt1;3;5;7* quadruple mutant exhibits a severe growth defect^44^, we used the *ipt3-2;5-1;7-1* triple mutant together with *cyp735a2-1* and *log7-1*. When five-day-old seedlings were transferred onto a medium containing 8 μM zeocin, root growth was less severely inhibited in the three mutants than in wild-type (Fig. 4a). Counting the cortical cell number in the MZ showed that after zeocin treatment for 24 h, the meristem size was reduced to 50% in wild-type, but to 96%, 78% and 79% in *ipt3-2;5-1;7-1*, *cyp735a2-1* and *log7-1*, respectively (Fig. 4b, c). These results suggest that enhanced production of cytokinin precursors by IPT3/5/(7) in shoots, and CYP735A2- and LOG7-mediated synthesis of iP and *t*Z in roots, are all involved in DSB-induced inhibition of root growth and reduction of the meristem size.

**Figure 4.**
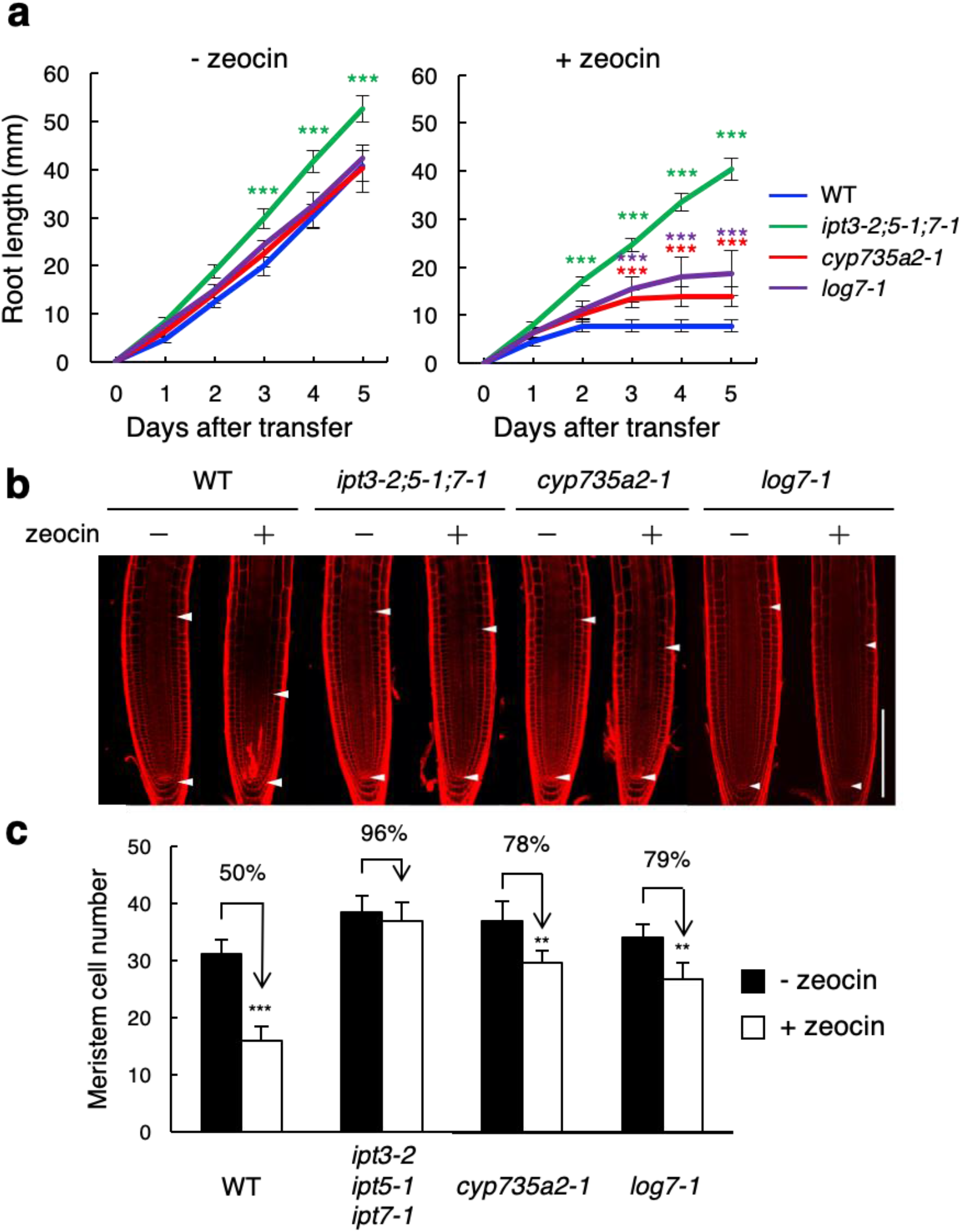
Roots of cytokinin biosynthesis mutants are tolerant to DSBs. (**a**) Root growth of wild-type, *ipt3-2;5-1;7-1*, *cyp735a2-1* and *log7-1*. Five-day-old seedlings were transferred to MS plates supplemented with (+ zeocin) or without (− zeocin) 8 μM zeocin, and root length was measured every 24 h. Data are presented as mean ± SD (n > 20). Significant differences from wild-type were determined by Student’s *t*-test: ***, *P* < 0.001. (**b**) Images of root tips. Five-day-old seedlings were treated with (+) or without (−) 8 μM zeocin for 24 h, and subjected to PI staining. Arrowheads indicate the QC and the boundary between the MZ and the TZ. Bar = 100 μm. (**c**) Cortical cell number in the MZ. Five-day-old seedlings were treated with (+ zeocin) or without (− zeocin) 8 μM zeocin for 24 h, and the number of cortical cells between the QC and the first elongated cell was counted. Data are presented as mean ± SD (n > 20). Significant differences from the control without zeocin treatment were determined by Student’s *t*-test: **, *P* < 0.01; ***, *P* < 0.001.

### Enhanced cytokinin signalling around the TZ is involved in DSB-induced meristem size reduction

It has been demonstrated that in *Arabidopsis* roots, cytokinin signalling is upregulated around the TZ due to localized expression of B-type ARRs, thereby restricting the meristem size^40^. Therefore, we speculated that a reduction of the meristem size under DNA damage conditions is a consequence of enhanced cytokinin signalling around the TZ, which is caused by higher cytokinin accumulation in the root tip. To test this possibility, we generated transgenic plants expressing *CYTOKININ OXIDASE 1* (*CKX1*), which encodes a cytokinin-degrading enzyme, in the TZ under the *RCH2* promoter (Supplementary Fig. 7a). When five-day-old seedlings were transferred onto zeocin-containing medium, root growth was only mildly inhibited in the two independent transgenic lines as compared with wild-type (Supplementary Fig. 7b). In the absence of zeocin, the cortical cell number in the MZ was higher in *pRCH2:CKX1* than in wild-type (*P* < 0.05, n = 20) (Supplementary Fig. 7c, d), matching a previous report^40^. Importantly, the transgenic lines displayed less reduction of the meristem size after zeocin treatment, implying lower sensitivity to DNA damage (Supplementary Fig. 7c, d).

Under normal growth conditions, one of the causes of meristem size restriction is ARR1/12-mediated induction of *SHY2*/*IAA3*, which encodes a negative regulator of auxin signalling, around the TZ^36^. As expected, the *SHY2* promoter activity was elevated around the TZ in response to zeocin treatment (Supplementary Fig. 8a). Moreover, the *shy2-31* loss-of-function mutant, which has a larger meristem than wild-type^36^, exhibited higher tolerance to DNA damage: zeocin-induced reduction of the meristem size was partially suppressed in *shy2-31* (*P* < 0.05, n = 20) (Supplementary Fig. 8b, c). This and the above results suggest that activation of cytokinin signalling around the TZ and a resultant increase of *SHY2* expression are associated with meristem size reduction after DNA damage.

### DSBs reduce auxin signalling in the root tip

SHY2 expressed around the TZ downregulates *PIN* expression, thereby inhibiting auxin transport and restricting the meristem size^36^. Therefore, the next question is whether auxin is associated with DDR in the root tip. We first examined the response of *PIN* genes to DNA damage. In roots, PIN1 and PIN4 are required for downward auxin flow in the stele, and PIN2 functions in upward transport in the lateral root cap and the epidermis^45^. PIN7 regulates downward auxin flow in the stele and redirection in the columella for lateral transport to the lateral root cap and the epidermis^45^. Our qRT-PCR data showed that the transcript levels of *PIN1* and *PIN4*, but not *PIN2* or *PIN7*, were reduced by 8 μM zeocin treatment (Fig. 5a). This result was supported by the analysis of *pPIN:PIN–GFP* reporter lines; in the absence of zeocin, PIN1–GFP and PIN4–GFP accumulate in the apical and basal parts of the vasculature, respectively^45^, whereas the GFP fluorescence was dramatically decreased by zeocin treatment (Fig. 5b, Supplementary Fig. 9). It is noteworthy that the PIN1–GFP signal diminished in the MZ although *SHY2* is induced in the TZ, but not in the MZ (Supplementary Fig. 8a). Since *PIN1* expression is upregulated by auxin in a time- and concentration-dependent manner^46^, it is likely that SHY2-mediated *PIN4* (and *PIN1*) repression around the TZ disturbs downward auxin flow, thereby reducing the auxin level and suppressing *PIN1* expression in the MZ. On the other hand, no change in expression patterns/levels of PIN2–GFP or PIN7–GFP was observed after zeocin treatment (Fig. 5b).

**Figure 5.**
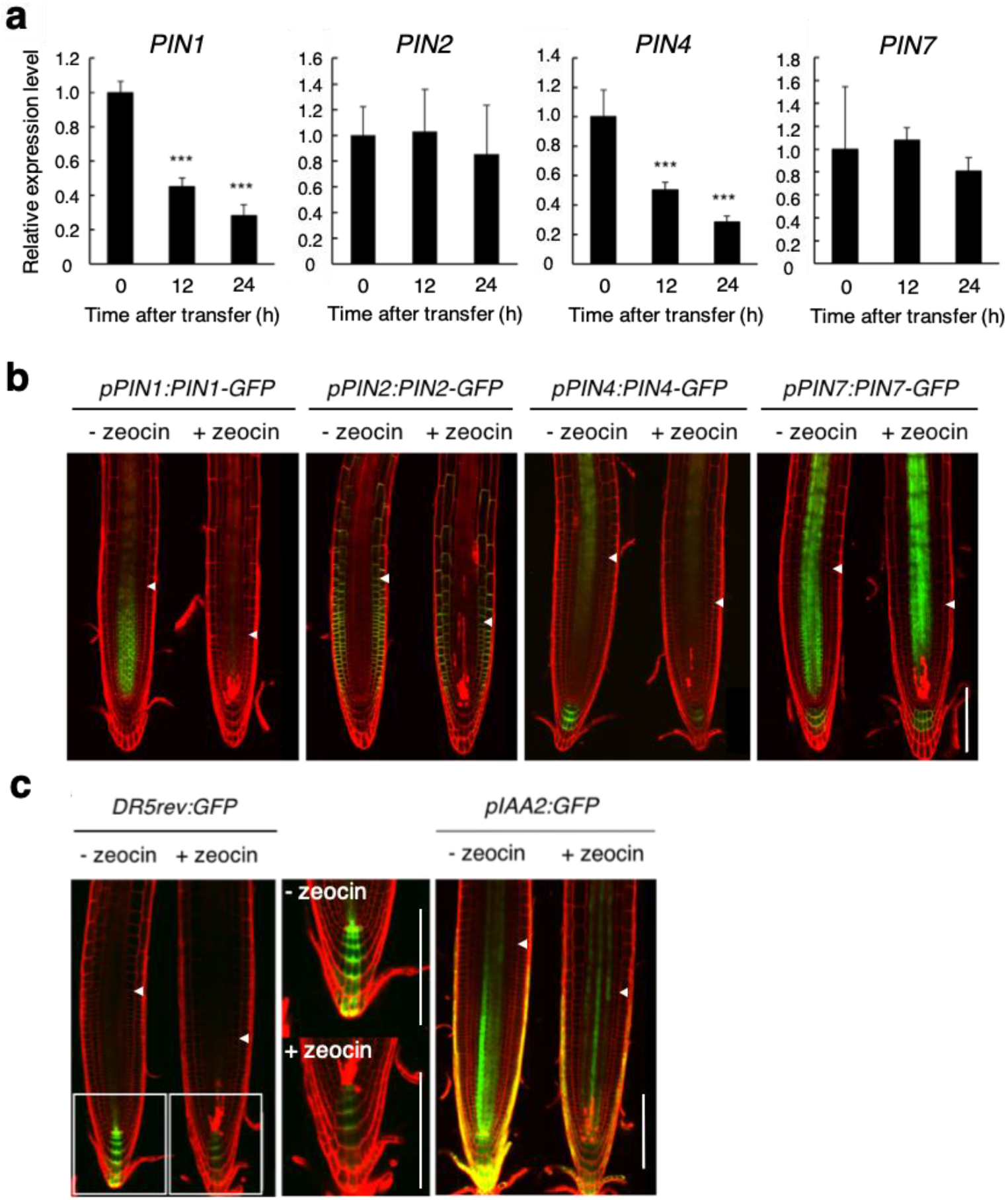
DSBs inhibit auxin transport and signalling in the root tip. (**a**) Transcript levels of *PIN1*, *PIN2*, *PIN4* and *PIN7*. Five-day-old wild-type seedlings were transferred to MS plates containing 8 μM zeocin, and grown for 0, 12 and 24 h. Total RNA was extracted from roots and subjected to qRT-PCR. Transcript levels of *PIN1*, *PIN2*, *PIN4* and *PIN7* were normalized to that of *ACTIN2*, and are indicated as relative values, with that for 0 h set to 1. Data are presented as mean ± SD calculated from three biological and technical replicates. Significant differences from the 0 h sample were determined by Student’s *t*-test: ***, *P* < 0.001. (**b**) Zeocin response of *pPIN1:PIN1-GFP*, *pPIN2:PIN2-GFP*, *pPIN4:PIN4-GFP* and *pPIN7:PIN7-GFP*. Five-day-old seedlings were treated with (+ zeocin) or without (− zeocin) 8 μM zeocin for 24 h. GFP fluorescence was observed after counterstaining with PI. Arrowheads indicate the boundary between the MZ and the TZ. Bar = 100 μm. (**c**) Zeocin response of *DR5rev:GFP* and *pIAA2:GFP*. Five-day-old seedlings were treated with (+ zeocin) or without (− zeocin) 8 μM zeocin for 24 h. GFP fluorescence was observed after counterstaining with PI. Magnified images of the areas marked by white boxes in *DR5rev:GFP* are shown on the right. Arrowheads indicate the boundary between the MZ and the TZ. Bars = 100 μm.

We then investigated the auxin response using the auxin-responsive marker *DR5:revGFP*^47^ and the GFP reporter driven by the *IAA2* promoter, which is one of the primary downstream targets of auxin signaling^47,48^. As shown in Fig. 5c, the *DR5:revGFP* signal in the QC and columella cells, and the *pIAA2:GFP* signal in the vasculature, QC and columella cells, were significantly reduced by 24-h zeocin treatment, indicating that DSBs decrease auxin signalling in the MZ, QC and the columella. Notably, zeocin-induced repression of *PIN1*, *PIN4* and *IAA2* was not observed in the *atm-2* or *sog1-1* mutant (Supplemental Fig. 10).

### Activation of cytokinin biosynthesis is crucial for reducing auxin signalling in the meristem

We then examined whether reduced auxin signaling in the MZ is caused by enhanced cytokinin biosynthesis. Our expression analyses showed that zeocin-triggered repression of *PIN1*, *PIN4* and *IAA2* was suppressed in the *log7-1* mutant (Fig. 6a). Moreover, PIN1–GFP accumulation was not decreased in *log7-1* after zeocin treatment (Fig. 6b). These results suggest that DSBs perturb downward auxin flow through activation of cytokinin biosynthesis.

**Figure 6.**
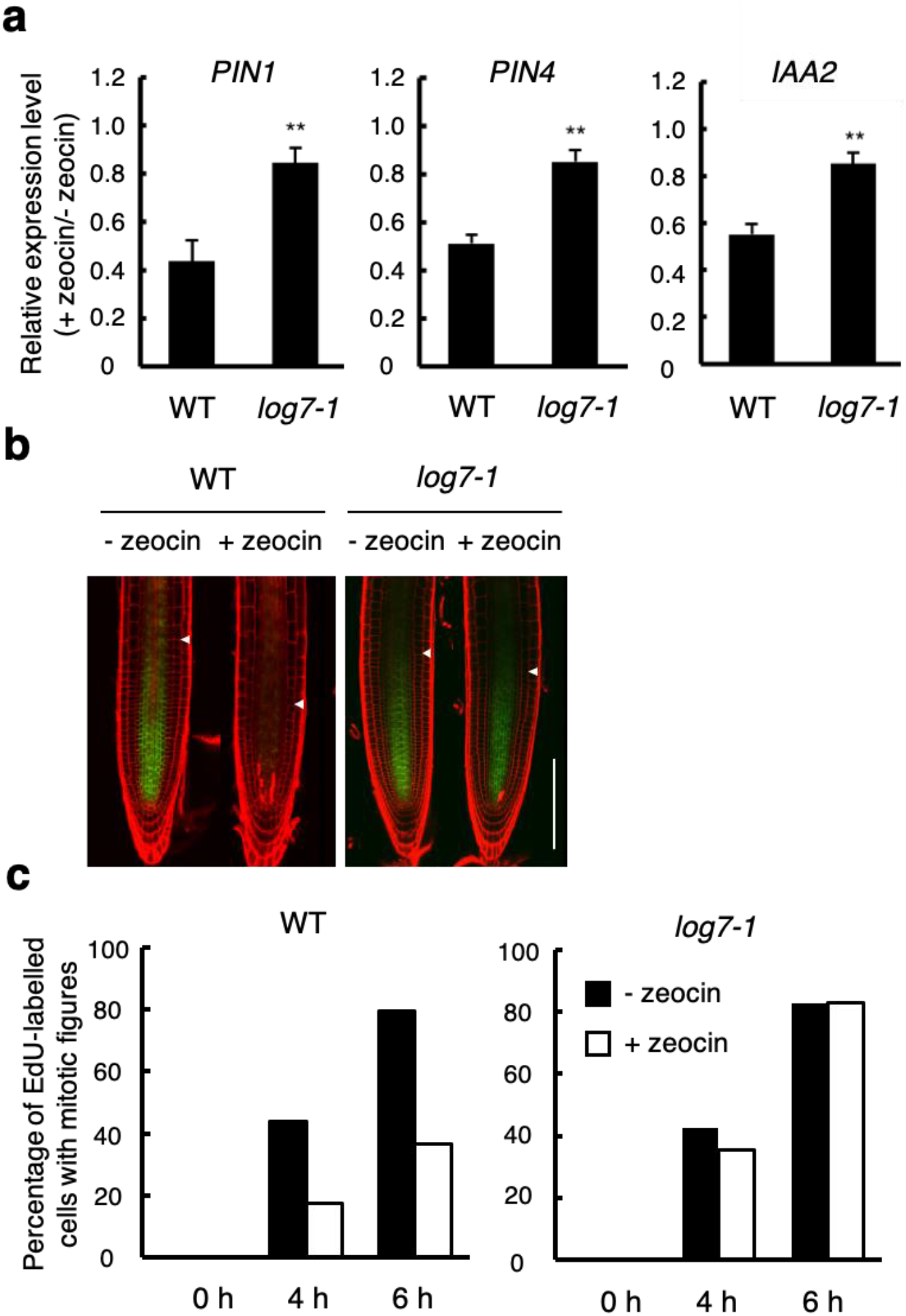
LOG7 is involved in DSB-induced suppression of auxin flow and cell division. (**a**) Transcript levels of *PIN1*, *PIN4* and *IAA2* in *log7*. Five-day-old wild-type (WT) and *log7-1* seedlings were transferred to MS plates with or without 8 μM zeocin, and grown for 24 h. Total RNA was extracted from roots and subjected to qRT-PCR. Transcript levels of *PIN1*, *PIN4* and *IAA2* were normalized to that of *ACTIN2*, and are indicated as relative values, with that for the control without zeocin treatment set to 1. Data are presented as mean ± SD calculated from three biological and technical replicates. Significant differences from wild-type were determined by Student’s *t*-test: **, *P* < 0.01. (**b**) *PIN1-GFP* expression in *log7*. Five-day-old seedlings of wild-type (WT) and *log7-1* harbouring *pPIN1:PIN1-GFP* were treated with (+ zeocin) or without (− zeocin) 8 μM zeocin for 24 h. GFP fluorescence was observed after counterstaining with PI. Arrowheads indicate the boundary between the MZ and the TZ. Bar = 100 μm. (**c**) G2 progression in *log7*. Five-day-old seedlings of wild-type (WT) and *log7-1* were transferred to MS plates with (+ zeocin) or without (− zeocin) 8 μM zeocin, and grown for 12 h. After pulse-labelling with 20 μM EdU for 15 min, seedlings were transferred back to MS plates with or without 8 μM zeocin, and grown for 0 h, 4 h and 6 h. Cells in the MZ were double-stained with EdU and DAPI, and those with mitotic figures were counted. Data are presented as the percentage of EdU-labelled cells among those with mitotic figures (n > 8).

### DSBs trigger G2 arrest by reducing auxin signalling

We previously reported that DSBs arrest the cell cycle preferentially at G2 in dividing cells^16^. To test whether enhanced cytokinin biosynthesis is also involved in G2 arrest in the MZ, we conducted EdU incorporation experiments to monitor cell cycle progression. EdU is incorporated into newly synthesized DNA during the S phase. After EdU-labelled cells pass through G2, cells which enter mitosis display mitotic figures^49^. Five-day-old wild-type and *log7-1* seedlings treated with or without 8 μM zeocin for 12 h were incubated with EdU for 15 min, and the number of EdU-labelled cells with mitotic figures in the MZ was counted after 4 h and 6 h. In wild-type, the percentage of these cells was significantly reduced in zeocin-treated roots, indicating retardation of G2 progression (Fig. 6c). However, in *log7-1*, we could not find any difference between zeocin-treated and non-treated samples (Fig. 6c). These data suggest that activation of cytokinin biosynthesis is associated with DSB-induced G2 arrest in the MZ.

We then asked whether reduced auxin signalling is the cause of DSB-induced G2 arrest. To address this issue, we treated wild-type roots with indole-3-acetic acid (IAA) at 5 nM, a very low concentration which did not change the meristem size at all (Fig. 7a, b). When 5 nM IAA was applied together with 8 μM zeocin, *LOG7* induction was observed similarly to the sole zeocin treatment (Fig. 7c), suggesting that 5 nM IAA does not affect DSB-dependent activation of cytokinin biosynthesis. However, IAA treatment partially suppressed zeocin-induced meristem size reduction (Fig. 7a, b). Moreover, EdU incorporation experiments showed that G2 arrest was also suppressed in the presence of both zeocin and IAA (Fig. 7d). These data suggest that a reduction in auxin signalling causes G2 arrest under DNA damage conditions.

**Figure 7.**
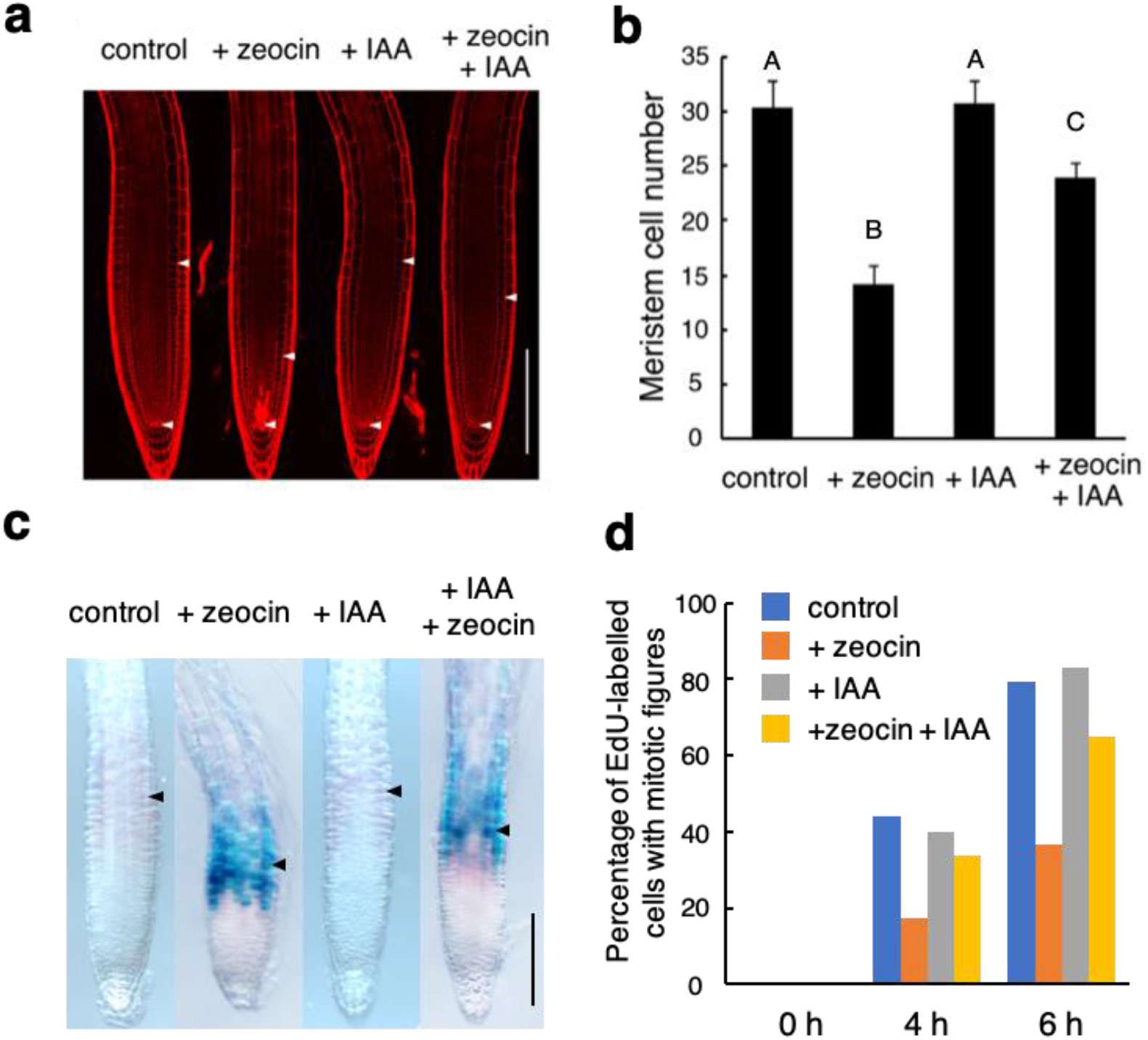
DSB-induced reduction in auxin signalling causes G2 arrest. (**a**) Root tips treated with zeocin and/or IAA. Five-day-old wild-type seedlings were transferred to MS plates supplemented with or without 8 μM zeocin and/or 5 nM IAA, and grown for 24 h, followed by PI staining. Arrowheads indicate the QC and the boundary between the MZ and the TZ. Bar = 100 μm. (**b**) Cortical cell number in the MZ treated with zeocin and/or IAA. Five-day-old wild-type seedlings were treated with or without 8 μM zeocin and/or 5 nM IAA for 24 h, and the number of cortical cells between the QC and the first elongated cell was counted. Data are presented as mean ± SD (n > 20). Bars with different letters are significantly different from each other (*P* < 0.01). (**c**) *LOG7* promoter activities after treatment with zeocin and/or IAA. Five-day-old *pLOG7:GUS* seedlings were transferred to MS plates supplemented with or without 8 μM zeocin and/or 5 nM IAA, and grown for 24 h, followed by GUS staining. Arrowheads indicate the boundary between the MZ and the TZ. Bar = 100 μm. (**d**) G2 progression in the presence of zeocin and/or IAA. Five-day-old wild-type seedlings were transferred to MS plates with or without 8 μM zeocin and/or 5 nM IAA, and grown for 12 h. After pulse-labelling with 20 μM EdU for 15 min, seedlings were transferred back to MS plates with or without 8 μM zeocin and/or 5 nM IAA, and grown for 0 h, 4 h and 6 h. Cells in the MZ were double-stained with EdU and DAPI, and those with mitotic figures were counted. Data are presented as the percentage of EdU-labelled cells among those with mitotic figures (n > 8).

### ARR2–CCS52A1 also functions in meristem size control in response to DNA damage

As mentioned above, the *shy2-31* mutation suppressed zeocin-induced meristem size reduction, but only partially (Supplemental Fig. 8b, c). This suggests that in addition to SHY2-mediated inhibition of auxin signalling, another mechanism functions in meristem size control. We previously reported that cytokinins promote an early onset of endoreplication in roots; in detail, cytokinin-activated ARR2, which is specifically expressed around the TZ, induces the expression of *CCS52A1*, encoding an activator of the E3 ubiquitin ligase APC/C, and promotes degradation of mitotic cyclins, thereby enhancing the transition from cell division to endoreplication^35^. Since cytokinins are elevated by DSBs as described above, it is likely that the ARR2–CCS52A1 pathway is also associated with meristem size reduction under DNA stress. To test this possibility, we first observed the promoter activity of *CCS52A1* using the GFP reporter line. As shown in Fig. 8a, the GFP signal in epidermal and cortical cells around the TZ was significantly elevated after zeocin treatment. qRT-PCR revealed that *CCS52A1* transcripts increased by about fourfold in the presence of zeocin, while no such induction was observed in *atm-2*, *sog1-1* or *log7-1* (Fig. 8b, c), suggesting that *CCS52A1* is upregulated through DNA damage signalling and by higher accumulation of cytokinins. Importantly, zeocin-induced meristem size reduction was less marked in the *ccs52a1-1* or *arr2-4* knockout mutant than in wild-type (Fig. 8d), suggesting that ARR2- and CCS52A1-mediated enhancement of endoreplication onset also contributes to meristem size reduction.

**Figure 8.**
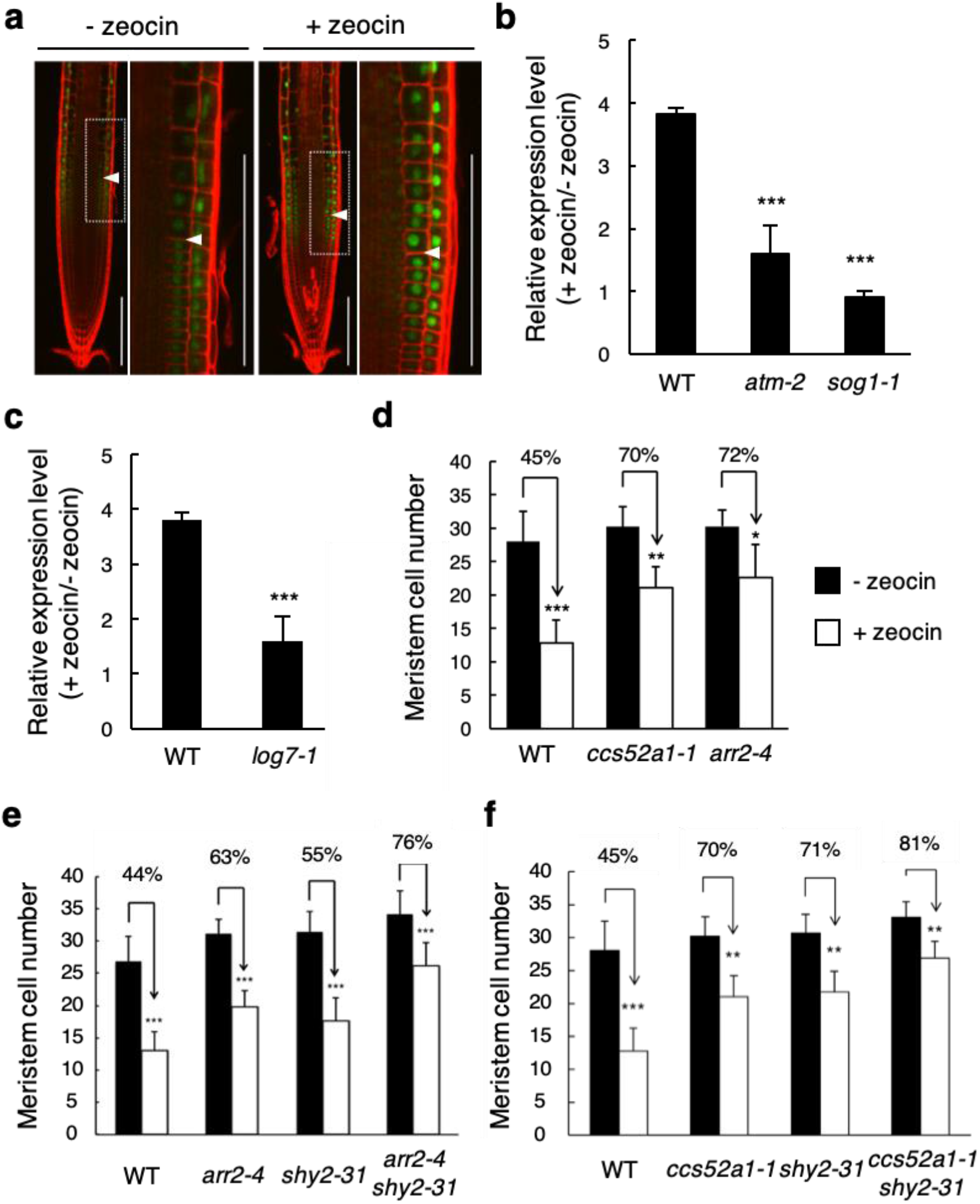
The ARR2–CCS52A1 pathway participates in meristem size reduction caused by DSBs. (**a**) *CCS52A1* expression in the presence of zeocin. Five-day-old *pCCS52A1:NLS-GFP* seedlings were transferred to MS plates with (+ zeocin) or without (− zeocin) 8 μM zeocin, and grown for 24 h. GFP fluorescence was observed after counterstaining with PI. Arrowheads indicate the boundary between the MZ and the TZ. Magnified images of the areas marked by white boxes are shown on the right. Bars = 100 μm. (**b, c**) Transcript levels of *CCS52A1* in *atm*, *sog1* and *log7*. Five-day-old seedlings were transferred to MS plates with or without 8 μM zeocin, and grown for 24 h. Total RNA was extracted from roots and subjected to qRT-PCR. Transcript levels of *CCS52A1* were normalized to that of *ACTIN2*, and are indicated as relative values, with that for the control without zeocin treatment set to 1. Data are presented as mean ± SD calculated from three biological and technical replicates. Significant differences from wild-type were determined by Student’s *t*-test: ***, *P* < 0.001. (**d–f**) Cortical cell number in the MZ. Five-day-old seedlings of wild-type (WT), *ccs52a1-1*, *arr2-4*, *shy2-31*, *arr2-4 shy2-31* and *ccs52a1-1 shy2-31* were transferred to MS plates with or without 8 μM zeocin, and grown for 24 h. The number of cortical cells between the QC and the first elongated cell was counted. Data are presented as mean ± SD (n > 20). Significant differences from the control without zeocin treatment were determined by Student’s *t*-test: *, *P* < 0.05; **, *P* < 0.01; ***, *P* < 0.001.

To reveal whether the SHY2-dependent signalling and the ARR2/CCS52A1-mediated pathway cooperate in DSB-induced meristem size reduction, we performed genetic experiments. After zeocin treatment, the root meristem size was reduced to 44% in wild-type, and to 63% and 55% in *arr2-4* and *shy2-31*, respectively, but only to 76% in the *arr2-4 shy2-31* double mutant (Fig. 8e). Similarly, the *ccs52a1-1 shy2-31* double mutant exhibited less sensitivity to zeocin as compared to each single mutant (Fig. 8f). These results suggest that both ARR1/12–SHY2 and ARR2–CCS52A1 pathways function in meristem size control under DNA stress.

## Discussion

In this study, we revealed that DSB-stimulated ATM and SOG1 enhance cytokinin biosynthesis. As a result, cytokinin signalling is activated around the TZ, and *SHY2* is upregulated to repress *PIN1* and *PIN4*, thereby perturbing downward auxin flow and causing cell cycle arrest in the meristem. At the same time, ARR2 induces *CCS52A1* and promotes an early onset of endoreplication. The combined effects of SHY2- and CCS52A1-dependent pathways play a crucial role in DNA damage-induced meristem size reduction (Fig. 9). It has been reported recently that cytokinin-activated ARR1 regulates root meristem size through the control of auxin quantity in the lateral root cap (LRC) by induction of GH3.17 and PIN5, which inactivate IAA and pump auxin from the cytoplasm into the endoplasmic reticulum, respectively^50^. In the present study, we have not investigated the involvement of GH3.17 and PIN5 in the DDR, but it is possible that enhanced cytokinin biosynthesis affects the auxin level in the LRC and reduces the meristem size. We previously demonstrated that DNA damage upregulates cytokinin signalling in the lateral root (LR) primordium and inhibits LR formation^51^. However, it has remained unknown whether the cytokinin level increases in the LR primordium and how LR development ceases after DNA damage. Since cytokinins are known to inhibit auxin-induced expression of *PIN*s during LR development^52^, it is probable that cytokinin signalling activated by DNA damage inhibits auxin flow by a mechanism similar to that shown in this study and disturbs proper organization of the LR meristem. Further studies will answer the question of whether the mechanisms identified here function in controlling growth of the whole root system under genotoxic stress conditions.

**Figure 9.**
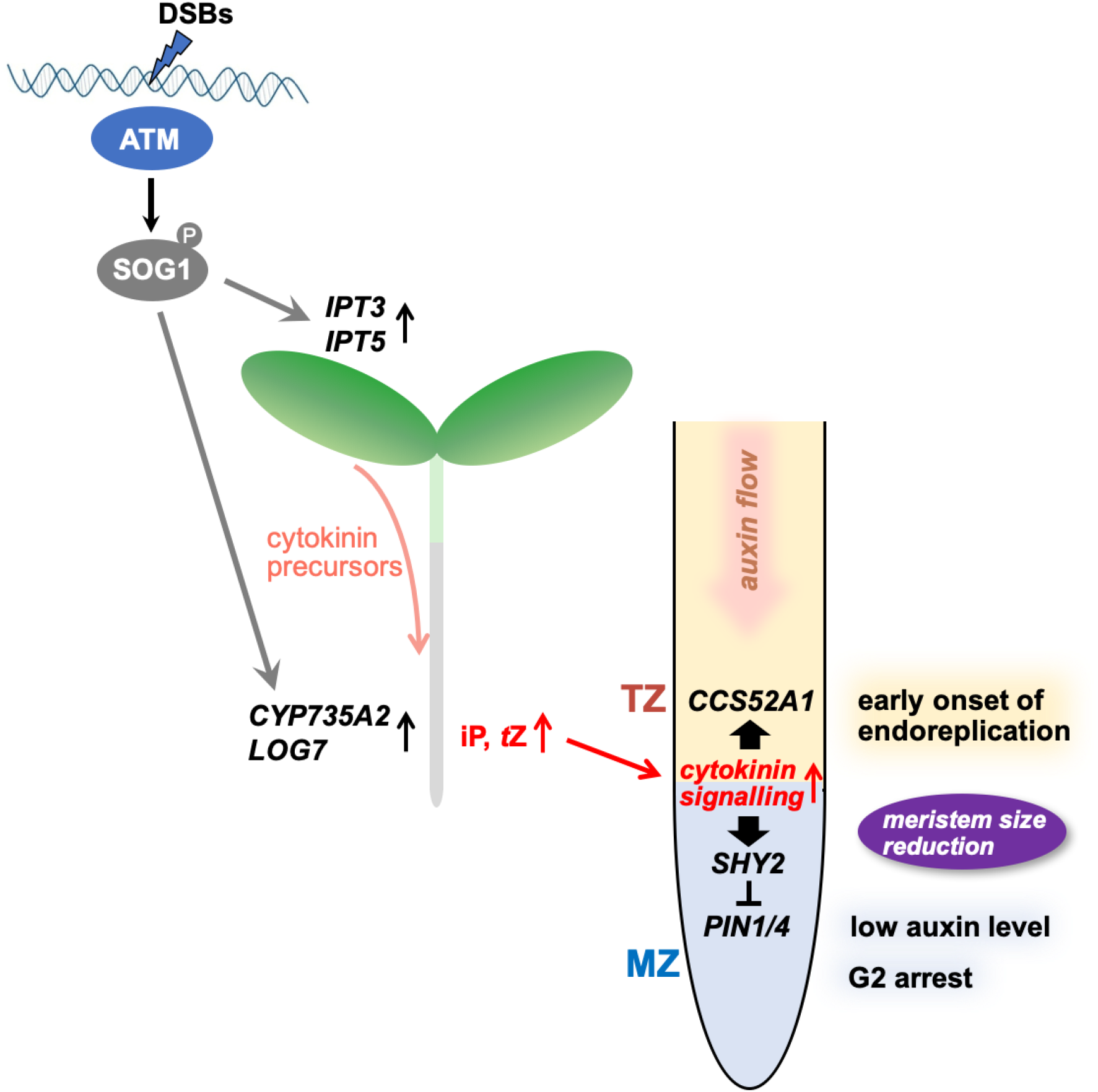
Model for DSB-induced reduction of the root meristem size. DSBs induce *IPT3* and *IPT5* in shoots and *CYP735A2* and *LOG7* in roots through the ATM– SOG1 pathway. As a result, iP and *t*Z levels are elevated in the root tip, and cytokinin signalling around the boundary between the MZ and the TZ is activated. This upregulates *SHY2* and suppresses *PIN1* and *PIN4* expression, thereby inhibiting downward auxin flow and reducing the auxin level in the MZ. Increased cytokinin signalling also upregulates *CCS52A1* around the boundary and promotes degradation of mitotic cyclins. These events cause G2 arrest in the MZ and an early onset of endoreplication, both of which trigger meristem size reduction and root growth inhibition.

One of the key findings in this study is that DNA damage induces the cytokinin biosynthesis genes *IPT1*, *IPT3*, *IPT5*, *IPT7*, *LOG7* and *CYP735A2*. Since these genes are not among the SOG1 target genes previously identified by ChIP-seq analyses^11,12^, downstream transcription factor(s) are likely involved in their induction. The *promoter:GUS* lines showed that *IPT1* expression increased in zeocin-treated roots, while *IPT3* and *IPT5* were induced in cotyledons and shoot apices, respectively (Fig. 3). Since the *ipt3-2;5-1;7-1* triple mutant exhibited a zeocin-tolerant phenotype in roots (Fig. 4), it is conceivable that cytokinin precursors produced by IPT3 and IPT5 are transported from shoots to roots and promote the DDR in the root tip. In support of this hypothesis, the levels of cytokinin precursors were significantly elevated in the root tip after zeocin treatment (Fig. 2). We could not detect an increase in the iP level, probably because iP is efficiently degraded by CKX enzymes and converted to *N*-glucosides^53,54^. Indeed, iP7G was highly increased by zeocin treatment in the TZ (Fig. 2). Interestingly, DSBs also induce *CYP735A2* and *LOG7* specifically around the TZ (Fig. 3), indicating that conversion from iP- to *t*Z-type cytokinins and the final activation step of cytokinin biosynthesis are enhanced around the TZ. It is noteworthy that LOG7 has the highest specificity constant (*K_cat_/K_m_*) among the eight LOG proteins in *Arabidopsis*^55^, suggesting that *LOG7* induction facilitates efficient production of biologically active cytokinins around the TZ, thereby enabling a rapid and strong response to DNA stress in terms of the control of meristem size and root growth.

We previously reported that Rep-MYBs, which repress the expression of G2/M-specific genes, play an essential role in inhibiting G2 progression in response to DNA damage^15^. Under normal growth conditions, CDKs phosphorylate Rep-MYBs and promote their degradation through the ubiquitin–proteasome pathway, leading to a release of transcriptional repression of G2/M-specific genes. In response to DNA damage, CDK activity is reduced, and Rep-MYBs are stabilized to cause G2 arrest^15^. This model indicates that how CDK activity is suppressed is a key to trigger G2 arrest when roots are exposed to DNA stress. The present study showed that DSBs activate cytokinin signalling around the TZ and inhibit downward auxin flow, thereby decreasing the auxin level in the meristem. Therefore, we propose that a decline in auxin level has a central role in reducing CDK activity in dividing cells. It has been shown that in *Arabidopsis*, auxin upregulates the expression of *CDKA;1*, which encodes the functional orthologue of yeast Cdc2/Cdc28p^56,57^. Genes for A2-type cyclins, and CDK inhibitors KIP-RELATED-PROTEIN 1 (KRP1) and KRP2, are induced and repressed, respectively, by exogenous auxin treatment^58,59^. Moreover, it was reported that treatment with the auxin antagonist α-(phenyl ethyl-2-one)-IAA rapidly decreased the transcript levels of core cell cycle regulators, such as CDKB1;1 and mitotic cyclin CYCA2;3^60^. These observations suggest that a decline in auxin level leads to a reduction in overall CDK activities, thereby enabling stabilization of Rep-MYBs. Indeed, Rep-MYBs are phosphorylated by all types of CDKs (A-, B1- and B2-types) in *Arabidopsis*^15^. Our recent study revealed that the transcription factors ANAC044 and ANAC085 are also involved in the control of Rep-MYB stability under DNA stress^17^; therefore, the next important question is how reduced auxin level and the ANAC044/085-dependent pathway cooperate in DNA damage-induced cell cycle arrest.

Previous reports demonstrated that various stresses reduce the cytokinin amount in shoots. For example, drought stress represses the expression of *IPT* genes and induces the *CKX* genes, thereby decreasing the cytokinin content in shoots^61^. Heat and salt stresses reduce endogenous cytokinin level by downregulating the *IPT* genes^61^. We also found that zeocin treatment suppressed *ARR5:GUS* expression in shoots, and that expression of *IPT9* and *LOG8* decreased in zeocin-treated leaves (Supplementary Fig. 11), suggesting distinct regulation of cytokinin level between shoots and roots. These observations are consistent with the role of cytokinins in cell cycle regulation, namely that cytokinins promote cell division in shoots but inhibit mitotic activity and promote endoreplication in roots^35,62^. We assume that stress signals control the cytokinin level differentially in shoots and roots, thus suppressing cell division and causing overall growth inhibition. Further studies will uncover the mechanisms underlying how cytokinin biosynthesis is controlled in distinct parts of plants in response to external stresses.

## Methods

### Plant growth conditions

*Arabidopsis thaliana* (ecotype Columbia-0) plants were grown vertically under continuous light conditions at 22 °C on Murashige and Skoog (MS) plates [0.5 × MS salts, 0.5 g/l 2-(*N*-morpholino)ethanesulfonic acid (MES), 1% sucrose and 1.2% phytoagar (pH 6.3)]. For DNA damage treatments, five-day-old seedlings were transferred to new MS plates supplemented with or without 8 μM zeocin (Invitrogen Life Technologies). For aluminium treatments, five-day-old seedlings were transferred to a medium [1 mM KNO_3_, 0.2 mM KH_2_PO_4_, 2 mM MgSO_4_, 0.25 mM (NH_4_)_2_SO_4_, 1 mM Ca(NO_3_)_2_, 1 mM CaSO_4_, 1 mM K_2_SO_4_, 1 μM MnSO_4_, 5 μM H_3_BO_3_, 0.05 μM CuSO_4_, 0.2 μM ZnSO_4_, 0.02 μM NaMoO_4_, 0.1 μM CaCl_2_, 0.001 μM CoCl_2_, 1% sucrose and 1.2% phytoagar (pH 4.2)]^10^ supplemented with or without 1.5 mM aluminium.

### Plant materials and constructs

*pARR5:GUS*^37^, *TCSn:GFP*^38^, *atm-2*^5^, *sog1-1*^8^, *pAHP6:GFP*^63^, *pAPL:GFPer*^64^, *pCO2:H2B-YFP*^65^, *pWOX5:NLS-YFP*^66^, *pSCR:GFP-SCR*^65^, *pRCH1:GFP*^40^, *pRCH2:CFP*^40^, *pIPT1:GUS*^43^, *pIPT3:GUS*^43^, *pIPT5:GUS*^43^, *pIPT7:GUS*^43^, *pCYP735A2:GUS*^23^, *pLOG7:GUS*^55^, *ipt3-2;5-1;7-1*^44^, *cyp735a1-2;a2-1*^23^, *log7-1*^55^, *pSHY2:GUS6*^67^, *pPIN1:PIN1-GFP*^68^, *pPIN2:PIN2-GFP*^68^, *pPIN4:PIN4-GFP*^46^, *pPIN7:PIN7-GFP*^68^, *DR5:revGFP*^47^, *pIAA2:GFP*^27^, *pCCS52A1:NLS-GFP*^35^, *ccs52a1-1*^35^, *arr2-4*^35^ and *shy2-31*^35^ were described previously. *sog1-1* (Col-0/L*er* background) was backcrossed to Col-0 wild-type three times before use.

The full-length open reading frame of *CKX1* was amplified from *Arabidopsis* cDNA by polymerase chain reaction (PCR) with F_CKX1 (5′-AAAAAGCAGGCTTCATGGGATTGACCTCATCCTTAC-3′) and R_CKX1 (5′-AGAAAGCTGGGTCTTATACAGTTCTAGGTTTCGGC-3′) primers, and cloned into the pDONR221 entry vector (Thermo Fisher Scientific) by a BP recombination reaction according to the manufacturer’s instructions. The *RCH2* promoter was amplified from *Arabidopsis* genomic DNA by PCR with FP_RCH2 (5′-ATAGAAAAGTTGGGAGGTAAGAATCATGAGAGTGGAG-3′) and RP_RCH2 (5′-TTGTACAAACTTGACATTTGCCTCAAAATACGAAAAGAAG-3′) primers, and cloned into the pDONRP4P1R entry vector (Thermo Fisher Scientific) by a BP recombination reaction. To generate the fusion construct of the *RCH2* promoter and *CKX1*, the entry clones were mixed with the R4pGWB601 destination vector^69^ in a LR recombination reaction according to the manufacturer’s instructions (Thermo Fisher Scientific). To generate the *pARR1:ARR1–GUS* and *pARR2:ARR2–GUS* constructs, the *ARR1* and *ARR2* genomic fragments were amplified from *Arabidopsis* genomic DNA by PCR with FG_ARR1 (5′-AAAAAGCAGGCTGACAACGATAGATGGAGAGG-3′) and RG_ARR1 (5′-AGAAAGCTGGGTAAACCGGAATGTTATCGATGG-3′) primers for *ARR1*, and FG_ARR2 (5′-AAAAAGCAGGCTGGATCCACGTATCAAGGGTC-3′) and RG_ARR2 (5′-AGAAAGCTGGGTAGACCTGGATATTATCGATGG-3′) primers for *ARR2*. Each PCR fragment was cloned into the pDONR221 entry vector by a BP recombination reaction, and subsequently transferred to the pGWB3 destination vector^70^ by a LR recombination reaction. All constructs were introduced into the *Agrobacterium tumefaciens* GV3101 strain harbouring the plasmid pMP90. The obtained strains were used to generate stably transformed plants by the floral dip transformation method.

### Quantitative RT-PCR

Total RNA was extracted from root tips or whole seedlings with the Plant Total RNA Mini Kit (Favorgen Biotech). First-strand cDNA was prepared from total RNA with the ReverTra Ace (Toyobo) according to the manufacturer’s instructions. For quantitative PCR, a THUNDERBIRD SYBR qPCR Mix (Toyobo) was used with 100 nM primers and first-strand cDNA. PCR reactions were run on a LightCycler 480 Real-Time PCR System (Roche) according to the following conditions: 95 °C for 5 min; 45 cycles at 95 °C for 10 s, 60 °C for 10 s, and 72 °C for 15 s. Transcript levels were normalized to that of *ACTIN2*. Three biological and technical replicates were performed for each experiment. The following primers were used: 5′-CTGGATCGGTGGTTCCATTC-3′ and 5′-CCTGGACCTGCCTCATCATAC-3′ for *ACTIN2*, 5′-AGAGATCACAACGAATCAGATTACGT-3′ and 5′-ATGACGCCGAGGAGATGGT-3′ for *IPT1*, 5′-CGGGTTCGTGTCTGAGAGAG-3′ and 5′-CTGACTTCCTCAACCATTCCA-3′ for *IPT3*, 5′-AGTTACAGCGATGACCACCA-3′ and 5′-GGCAGAGATCTCCGGTAGG-3′ for *IPT5*, 5′-ACTCCTTTGTCTCAAAACGTGTC-3′ and 5′-TGAACACTTCTCTTACTTCTTCGAGT-3′ for *IPT7*, 5′-CATGTTCTAGGGGTCATTCCA-3′ and 5′-CTCCGATGGTCTCACCAGTT-3′ for *CYP735A2*, 5′-TGATGCTTTTATTGCCTTACCA-3′ and 5′-CCACCGGCTTGTCATGTAT-3′ for *LOG3*, 5′-GTTTGATGGGTTTGGTTTCG-3′ and 5′-CACCGGTCAACTCTCTAGGC-3′ for *LOG4*, 5′-CTCCGATGGTCTCACCAGTT-3′ and 5′-CATGTTCTAGGGGTCATTCCA-3′ for *LOG7*, 5′-ATTGCACTCCCTGGAGGTTA-3′ and 5′-CCCATCAACATTCAATAGACCA-3′ for *LOG8*, 5′-CTCTCCACGTACTGGTTGTCGTTAC-3′ and 5′-CGGAAGAGATCTTCACATGTTTGTG-3′ for *PIN1*, 5’-AAGTCACGTACATGCATGTG-3’ and 5’-AGATGCCAACGATAATGAGTG-3’ for *PIN2*, 5′-CGAAAGAGTAATGCTAGAGGTGGTG-3′ and 5′-AATATCAGTCGTGTCATCACACTTG-3′ for *PIN4*, 5′-CGAAAGAGTAATGCTAGAGGTGGTG-3′ and 5′-AATATCAGTCGTGTCATCACACTTG-3′ for *PIN7*, 5’-GAAGAATCTACACCTCCTACCAAAA-3’ and 5’-CACGTAGCTCACACTGTTGTTG-3’ for *IAA2*, and 5′-CACGCTGCAAGAGAACAAGA-3′ and 5′-ACCACTTGAGTCCGCATACC-3′ for *CCS52A1*.

### GUS staining

Seedlings were incubated in a GUS staining solution [100 mM sodium phosphate, 1 mg/ml 5-bromo-4-chloro-3-indolyl ß-D-glucuronide, 0.5 mM ferricyanide, and 0.5 mM ferrocyanide (pH 7.4)] in the dark at 37 °C. The samples were cleared with a transparent solution [chloral hydrate, glycerol and water (8 g:1 ml:1 ml)], and observed under a light microscope (Olympus).

### Microscopic observation

Five-day-old roots were stained with 10 μM propidium iodide solution for 1 min at room temperature, and root tips were observed under a confocal laser scanning microscope (Olympus, FluoView FV1000).

### Root growth analysis

Seedlings were grown vertically in square plates, and root tips were marked every 24 h. Plates were photographed, and root growth was calculated by measuring the distance between successive marks along the root axis with ImageJ software (http://rsb.info.nih.gov/ij/).

### Quantification of cytokinin levels in isolated cell populations

Protoplasts were isolated from root tips of five-day-old *pRCH1:GFP* and *pRCH2:CFP* seedlings, which were treated with or without 8 μM zeocin for 24 h. GFP- or CFP-positive protoplasts were collected through FACS, and analysed for their cytokinin concentration using liquid chromatography–tandem mass spectrometry (LC–MS/MS). Protoplast isolation, cell sorting, cytokinin purification and quantification were performed as described previously^41^. Analysis and sorting were performed using BD FACSAria I and BD FACSDIVA software. GFP fluorescence was excited using a 488-nm laser and collected using a FITC filter set (emission filter: 530 ± 30 nm), while CFP fluorescence was excited using a 405-nm laser and collected using an Alexa Fluor 430 filter set (emission filter: 550 ± 30 nm). The collected fluorescent protoplasts were snap-frozen in liquid nitrogen after each sorting experiment. Prior to cytokinin purification and subsequent quantification, the samples were thawed on ice and grouped in pools to form four biological replicates per genotype and treatment (60,000–160,000 protoplasts per replicate).

### EdU-pulse labelling

Five-day-old seedlings were grown on MS medium supplemented with or without 8 μM zeocin for 12 h, and transferred to MS medium with or without zeocin and with 20 μM EdU for 15 min. After washing with MS medium, seedlings were transferred back to MS medium supplemented with or without zeocin. EdU staining was performed with a Click-iT Plus EdU Alexa Fluor 488 Imaging Kit (Thermo Fisher Scientific) according to the manufacturer’s instructions. Roots were double-stained with EdU and DAPI, and observed with a confocal microscope (FV1000, Olympus).

### Data availability

Data supporting the findings of this work are available within the paper and its Supplementary Information files. A reporting summary for this article is available as a Supplementary Information file. The source data are provided with this paper.

## Acknowledgements

We thank Tatsuo Kakimoto, Anne Britt, Yoshikatsu Matsubayashi, Mitsuhiro Aida, Ildoo Hwang, Tsuyoshi Nakagawa, Shunsuke Miyashima, Keiji Nakajima, and the *Arabidopsis* Biological Research Center for providing *Arabidopsis* mutants, binary vectors, and transgenic plants. We also thank Mikiko Kojima for her help in cytokinin measurements. This work was supported by MEXT KAKENHI (Grant Numbers 22119009, 26113515, 17H06470 and 17H06477), JSPS KAKENHI (Grant Numbers 26291061, 26650099, 26840096 and 19K06708), and JST CREST. KL and IA were supported by the Knut and Alice Wallenberg Foundation, the Swedish Research Council and the Swedish Governmental Agency for Innovation Systems.

## Contributions

N.T., S.I., K.N., I.A., M.K. and K.L. performed experiments. N.T., S.I., K.N., I.A., M.K. and K.L. performed data analysis. N.T., S.I., H.S., I.A., K.L. and M.U. designed experiments. N.T. and M.U. wrote the manuscript.

## Competing financial interests

The authors declare no competing financial interests.

